# A solinvivirus induces cellular antiviral response and reduces the lifespan of adult *Hermetia illucens* (black soldier flies)

**DOI:** 10.1101/2025.09.19.677374

**Authors:** Robert D. Pienaar, Pablo García-Castillo, Harmony Piterois, Violette Wallart, Frédéric Manas, Elisabeth A. Herniou, Salvador Herrero

## Abstract

Viral pathogens pose an emerging threat to the sustainability of insect mass-rearing systems, yet remain understudied in key species like the black soldier fly (BSF, *Hermetia illucens*). Although multiple viral sequences have been reported in BSF, their role in disease has not been established until now. Here, we provide the first *in vivo* characterisation of *Hermetia illucens* solinvivirus (HiSvV), confirming its role as a viral entomopathogen of BSF. Metatranscriptomic analysis of a diseased colony revealed a high viral load attributable to HiSvV. We successfully isolated the virus and developed injection- and oral-based infection assays to investigate replication, tissue tropism, transmission and risk of mortality. HiSvV replicated in inoculated adults, induced premature mortality in flies, and was transmitted both horizontally and vertically. Infected flies also mounted a broad antiviral response, supporting active pathogenesis. These findings establish HiSvV as the first confirmed viral pathogen of BSF and underscore the urgent need for viral surveillance and experimental tools to safeguard industrial insect rearing.

## Introduction

Black Soldier Flies (BSF), *Hermetia illucens*, are increasingly used in waste management and as a source of protein in animal feed, making its health and productivity crucial for these industries [1–5]. Studies on entomopathogens affecting BSF are limited, presenting a gap in BSF pathology that has been highlighted on multiple occasions [4,6–9]. Although there has been a growing number of studies on the effect of more general insect pathogens [10–14], the first natural pathogen of BSF was only recently described in 2023 [8]. The authors described a *Paenibacillus* sp.-related pathogen causing the death of BSF with a syndrome previously described as “soft rot”.

Issues in insect mass-rearing can also arise from a less conspicuous agent, such as viruses [15,16]. Often, the presence of viruses can be missed, particularly since signs can mimic those of sub-optimal rearing practices, such as smaller larvae, lower feeding rates, and reduced fecundity [15–17]. In other cases, viral infections may remain unnoticeable until certain factors trigger the infection to produce obvious symptoms including bloated individuals, even death [16,18–20]. Since 2022, eight exogenous viruses associated with BSF have been discovered [7,9,21,22]. Among them, four were classified into the families *Dicistroviridae, Iflaviridae, Rhabdoviridae* and *Solinviviridae* and are closely related to other known insect pathogens [16,23–26]. However, no additional evidence was obtained on the potential pathogenicity of these viruses.

*Solinviviridae* is a recently described viral family of the *Picornavirales* with non-segmented, linear, positive-sense RNA genomes of approximately 10-11 kb, suspected of including an array of arthropod pathogens [25]. According to the ICTV, [25], there are about 69 viral sequences closely related to *Solinviviridae*, but only two species are accepted within the family. Despite this high number, few studies describe the actual viruses or their interactions with their hosts [27–30]. The most well described viruses in the *Solinviviridae* family are *Solenopsis invicta* virus 3 (SINV3), *Nylanderia fulva* virus 1 (NfV1), and two closely-related viruses, *Apis mellifera* solinvivirus 1 (AmSV1) and *Penaeus vannamei* solinvivirus (PvSV), which are not yet formally accepted as *Solinviviridae*. For SINV3, high virus titre has been linked to mortality and can cause colony collapse and decreased fecundity in ants [27,31,32]. The ecology of the *Solinviviridae* is not very well established, since the bulk of the family characterization work has been done using mainly SINV3 and NfV1 [25,27,28,30,31,33–41]. A common feature in

*Picornavirales* (including *Iflaviridae* and *Dicistroviridae*) closely related to *Solinviviridae* is that infections in insect rearings can go unnoticed until an outbreak occurs, often resulting in a strong display of symptoms and colony collapse [15,18,31,42,43].

The goal of this study was to unravel the potential pathogenic effect of viruses in BSF adults derived from a BSF rearing facility experiencing unexplained mortalities. After identifying HiSvV as the primary candidate responsible of the mortality experienced in a BSF colony, the virus was isolated and its interactions with BSF further characterised. To do that, we studied HiSvV transmission, pathology, and the transcriptional response of BSF to the infection. This comprehensive approach allowed us to better understand HiSvV biology and assess its risk and relevance to the BSF mass-rearing industry.

## Methods and materials

### BSF laboratory colony

A starter colony was provided by Entomotech S.L (Almeria, Spain) and the rearing was originally installed at the CBP research group at University of Valencia (Spain) in 2022. This colony was considered as “virus free” after initial screenings to use for HiSvV infection studies [22]. The individuals were reared at 28 °C on Gainesville diet (50% Wheat germ, 30% Alfalfa and 20% corn flour) in a similar manner to Deruytter *et al*., [44], but adding 150% MilliQ water to dry diet mix (0.3 kg of diet to 0.45 L of water). Essentially, 0.3 L (100 %) of water was initially added to 0.3 kg diet and was kept at 4 °C overnight to allow the diet to absorb the moisture. Before the diet was provided to the neonates or larvae, the remaining 0.15 L (50%) of water was mixed into the premoistened diet. Eggs laid were collected and incubated in parafilm sealed petri dishes, until the neonates hatched and were then placed onto 60g of diet into 0.5 L container. Five- to seven-days-old larvae were then moved onto 250 g of fresh diet within a 2 L container until pupation. The prepupae and pupae were manually sorted from the substrate and placed into the 2 L containers with a dry sheet of paper towel until emergence. Adults were then placed in cages (47.5 cm x 47.5 cm x 47.5 cm) and a cotton ball soaked in MilliQ water was maintained every two days.

### Virus detection in transcriptomic data

Reads from two metatranscriptomic datasets (NCBI sequence read archives SRR28203243 and SRR28203244) obtained from BSF adults originally received from a BSF rearing facility experiencing unexplained mortalities, where analysed for the presence of viruses infecting *Hermetia illucens*, as done by Pienaar *et al*., [22]. Through this approach, Krona charts generated using Lazypipe 2 were used to visualize viral diversity within the metatranscriptomic datasets relative to the BSF reference genome (GCF_905115235.1) [45,46]. Additionally, raw reads were trimmed using fastp v0.23.2 [47] and mapped to the reference genome of HiSvV (PQ228193). Bowtie2 v2.4.2 [48] and samtools v1.9 [49] were used to map and filter the unmapped reads from the BAM files. The mean coverage was then assessed using Geneious Prime v2021.1 (https://www.geneious.com) and the coverage plots were generated using the same script as previously described [22].

### Virus partial purification and TEM

Pools of BSF individuals (either pupae or adults) were homogenised inside a 1.5 ml centrifuge tube or 50 ml falcon tube, depending on the number of individuals. Then virus particles were partially purified using the virus prepurification protocol used by Hernández-Pelegrín et al., [50], however, centrifugations were performed at 10 000x *g*. When needed, a concentration step was performed by 10% PEG 6000 precipitation, and the pellet was resuspended in 100 µl of 1x PBS.

For transmission electron microscopy (TEM), prepurified virus (PPV) underwent negative staining using 2% phosphotungstate (PTA) after samples were fixed to a carbon-coated grid. The grids were visualized and captured using an HT7800 *RuliTEM* 120 kV transmission electron microscope (Hitachi, Chiyoda City, Japan). The mean size of the viral-like particles was measured for 30 viral particles using ImageJ v1.54g [51].

### Virus detection by quantitative RT-PCR (RT-qPCR)

RNA extractions were performed as in Pienaar *et al*., [22]. Post extraction, a DNase treatment was performed on the RNA using the DNase I, RNase-free kit (EN0521, Thermo Fisher Scientific, Waltham, MA, USA), followed by cDNA synthesis using the PrimeScript RT Reagent Kit (Perfect Real Time) (TAKRR037A, Takara Bio, Kusatsu, Japan). The reverse transcription quantitative PCRs (RT-qPCRs) were performed as in Pienaar *et al*., [22]. The relative abundance of the viral targets was obtained using the *RPL*8 housekeeping gene as described in Herrero *et al*., [52]. The plots were generated using the relative abundance values and ggplot2 v3.5.0 [53] in R v4.2.2 [54]. The quantification of isolated virus was measured as “genomic equivalents”, calculated using “Amplification factor^(Y intercept - Ct)^”.

### Viral infections by injection of prepupae and pipette-feeding of adults

The effect of infection on the colony was initially assessed by viral injection during the prepupal stages. For that, four experiments were performed on different generations using 100 prepupae for each treatment. Prepupae were sorted from substrate and washed three times with MilliQ water and placed on paper towel to dry before injection. For inoculation, the needle was inserted dorsolaterally by the end at an angle close to 90° into the tegument connecting the second and third segments from the posterior end (Figure S1A).

Individuals were injected with 5 µl of either 1x PBS solution or PPV containing 4 x 10^8^ HiSvV genomic equivalents/µl. Post-injection, the prepupae were placed into 2L pupation boxes but without any substrate and left to pupate at 25 °C with a lighting routine set to 12 hr:12 hr. From nine days post injection, the containers were checked daily for emerged adults. Once emerged, to collect adults, the containers were placed at 5 °C for a maximum of 10 minutes and the adult males and females were placed separately into fresh containers precooled on ice for counting and sex identification. Afterwards, they were placed at 28 °C for further experiments and were maintained in complete darkness except adults selected for cohabitation experiments. The rate of emergence and sex was recorded and visualized using ggplot2 and a Kruskal-Wallis rank sum test [55] in R stats v4.2.2 [54] was used to test the overall statistical differences between emergences.

Adults were inoculated by pipette-feeding them a solution containing prepurified HiSvV. To do that, adults were collected within 24 hours post-emergence and then males and females were retained in two separate dehydration boxes overnight (Figure S1B). Each dehydration box would contain a 1 to 2 cm deep layer of wood shavings. For the inoculation, individuals were held inside a P20 pipette tip by gently applying pressure to their thorax against the lip of the tip opening with their wings outside and allowed to drink (Figure S1C). The abdomen windows were observed for visible dye to confirm that the flies were drinking (Figure S1D). The inoculation solution was preprepared as follows per 4 µl dose: 1 µl 5% sucrose water, 1 µl of food colouring and 2 µl HiSvV PPV or 2 µl 1x PBS.

Each droplet-feeding experiment consisted of two treatments, HiSvV inoculated and mock infection (PBS control). Inoculated adults were placed individually in 120 ml cups and 1 ml of MilliQ water was provided directly to a cotton ball placed in between the cup lid a piece of paper towel (Figure S1B). The cups were stored in an incubator at 25 °C with a 12 hr:12 hr (light:dark) lighting regime.

### Assessment of HiSvV replication in BSF adults

The replication of HiSvV was monitored in adults for both the prepupal injection assays and for oral injection assays. For prepupae inoculated by injection, adult males and females were collected separately at various time points post-emergence (pe). Additionally, adults that died during cohabitation between 8–13 days post-emergence (dpe) were collected. In each case, pools of two individuals were prepared.

For orally infected adults, two males and two females were collected at different time points post-inoculation for qPCR as described above.

### Tropism of HiSvV in adult BSF

Virus tropism was determined in adults derived from the injected prepupae (3 out of four experiments). From each experiment, one male and female were collected at three days post emergence. Adults were washed with TE buffer (10mM Tris-HCl pH 8-8.6, 1mM EDTA) beforehand, then the head, wings and legs were dissected and placed into tubes according to body parts. Afterwards, the corpse was submerged into TE buffer and from the abdomen, the reproductive organs were removed, followed by the fat bodies, mid- and hindgut with Malpighian tubules, and lastly the thoracic muscles. Each piece of anatomy was rinsed three times in TE buffer before transfer to 300 µl of TRItidy G reagent (AppliChem GmbH, Darmstadt, Germany).

### HiSvV transmission

Cohabitation experiments were designed in order to evaluate the horizontal and vertical transmission of the virus. To implement that, fifteen individuals of each sex were selected to place 30 adults in a “25 cm x 25 cm x 25 cm” cage. For each experiment four cages were setup, each with a different combination of males and females. The following combinations used were virus positive males and females, virus negative males and females, virus positive males with virus negative females, and finally, virus negative males with virus positive females. This step was repeated four times using individuals from different generations. Prepupal-injected males and females emerging from pupae as adults within 24-48 hours of each other were used for each experiment and adults inside cages were maintained in a 12-hour light and 12-hour darkness regime. The cages were checked daily for eggs and dead adults, which were subsequently stored at -80 °C to test for horizontal (adults) and vertical (eggs) transmission of HiSvV using RT-qPCR as described above.

### Adult BSF survival bioassays after HiSvV infection

Adults were orally infected as described above and their mortality was monitored. The adults were incubated at 25°C for up to 30 days and 1 ml of MilliQ water was provided daily. The experiments were performed using HiSvV concentrations of 1.6×10^1^ and 1.6×10^4^ genome equivalents/µl, negative controls were mock infected with PBS. Thirty females and 30 males were used for each treatment. Mortality was recorded daily, and survival curves were plotted using survfit from the survival v3.5.8 R package [56,57] and ggsurvplot from the ggplot2 R package. Type II ANOVAs from the car v3.1.2 R package [58] were then performed on Cox proportional hazard models (survival R package) to test factors such as sex and experiment.

### Differential gene expression analysis

The impact of viral infection on BSF gene expression was analysed by RNAseq. To do that, RNA was extracted from adults orally infected with HiSvV at different time points. Library preparation and high-throughput sequencing was carried out as previously described in Pienaar *et al*., [22]. Datasets can be found at NCBI (bioproject PRJNA1079553). To obtain BSF transcripts, reads from long non-coding RNA (LncRNA) and messenger RNA (mRNA) datasets were trimmed, and poor-quality reads were removed using fastp. Then the reads were mapped to the BSF reference genome (GCF_905115235.1) using HISAT2 v2.2.1 [59]. Using samtools, the mapped reads were sorted, filtered and SAM files were converted to BAM files. An R script from Cerqueira de Araujo et al., [60] utilizing featureCounts from the Rsubread R package v2.8.2 [61] was used in combination with a GTF file for the BSF reference genome to obtain a gene count table from the BAM files. The DESeq2 R package v1.38.3 [62] was then used to contrast treatments HiSvV and mock (PBS) and obtain the log 2-fold changes (L2FCs) of genes using the design model ∼ “Treatment + Sex”. The gene counts for the whole transcriptome were normalized using DESeq and the counts of genes with a significant L2FC (*padj* < 0.05) were extracted. This was then repeated for males and females separately only using treatment as a factor in the design model. To visualise the significant up- and down-regulated differentially expressed genes (DEGs), the DESeq results were plotted in the form of volcano plots using R. The EggNOG mapper webserver [63,64]; accessed 29^th^ February 2024) was used to annotated functional and protein family (PFAM) descriptions for the protein sequences related to the BSF reference genome. From this, the significant DEGs were labelled with their corresponding EggNOG descriptions and PFAM. STRING [65] was then used to annotate the refseq BSF proteome (GCF_905115235.1; string taxid STRG0A70YGC) using the STRING database v12.0. Following, a protein interaction network was constructed for the male and female related DEG protein sequences using the STRING multiple protein search option with the default settings (STRG0A70YGC; accessed 30^th^ April 2025). The STRING protein search also included a gene ontology (GO) enrichment analysis, as well as KEGG and Reactome pathway searches for the DEG proteins. The protein sequences with biological process (BP) GO terms related to immunity were highlighted and the KEGG and Reactome pathway descriptions assigned to the related DEGs were extracted. Additionally, the relation to immunity for extra DEG was putatively determined through text-mining of the gene/protein names and a literature search. Lastly, the normalized counts from the male and female DESeq results were log2 transformed and the transformed counts for the DEGs with immune-related BP terms were extracted and used to construct a heatmap with row scaled Z-scores plotted, using pheatmap v1.0.12 [66].

### Viral small RNA profiling

The small RNA sequencing datasets were trimmed and mapped to HiSvV and processed using the same approach as above. Using sRNAplot [67] to generated sRNA profiles from the mapped and filtered bam files, the script was modified to also produce a csv datafile output. An R script then used the csv file to generate the profile plots and collations.

## Results

### High abundance of HiSvV reads in the diseased colony

A deep-sequencing shotgun transcriptomic approach was employed to analyse the presence of virus-related reads in a BSF colony that had high-levels of premature adult mortality. Two samples were analysed: one asymptomatic and freshly emerged adult fly (HA; healthy adult) and one adult fly that had died prematurely (DA, diseased adult). In the transcript data of the BSF adults, other than bacteriophage reads, only viral reads related to *Solinviviridae* were detected in both the HA and DA samples, although with striking difference in abundances (Figure 1A). When mapping the reads to the reference sequence for HiSvV (PQ228193), the profiles were similar for both datasets and spanned the entire reference sequence although the coverage was 500 times higher in diseased samples (coverage in DA = 147 343x and in HA=296x; Figure 1B). This higher viral abundance in the sample derived from the dead adult suggests HiSvV as responsible of the premature mortality observed in some adults.

**Figure 1.**
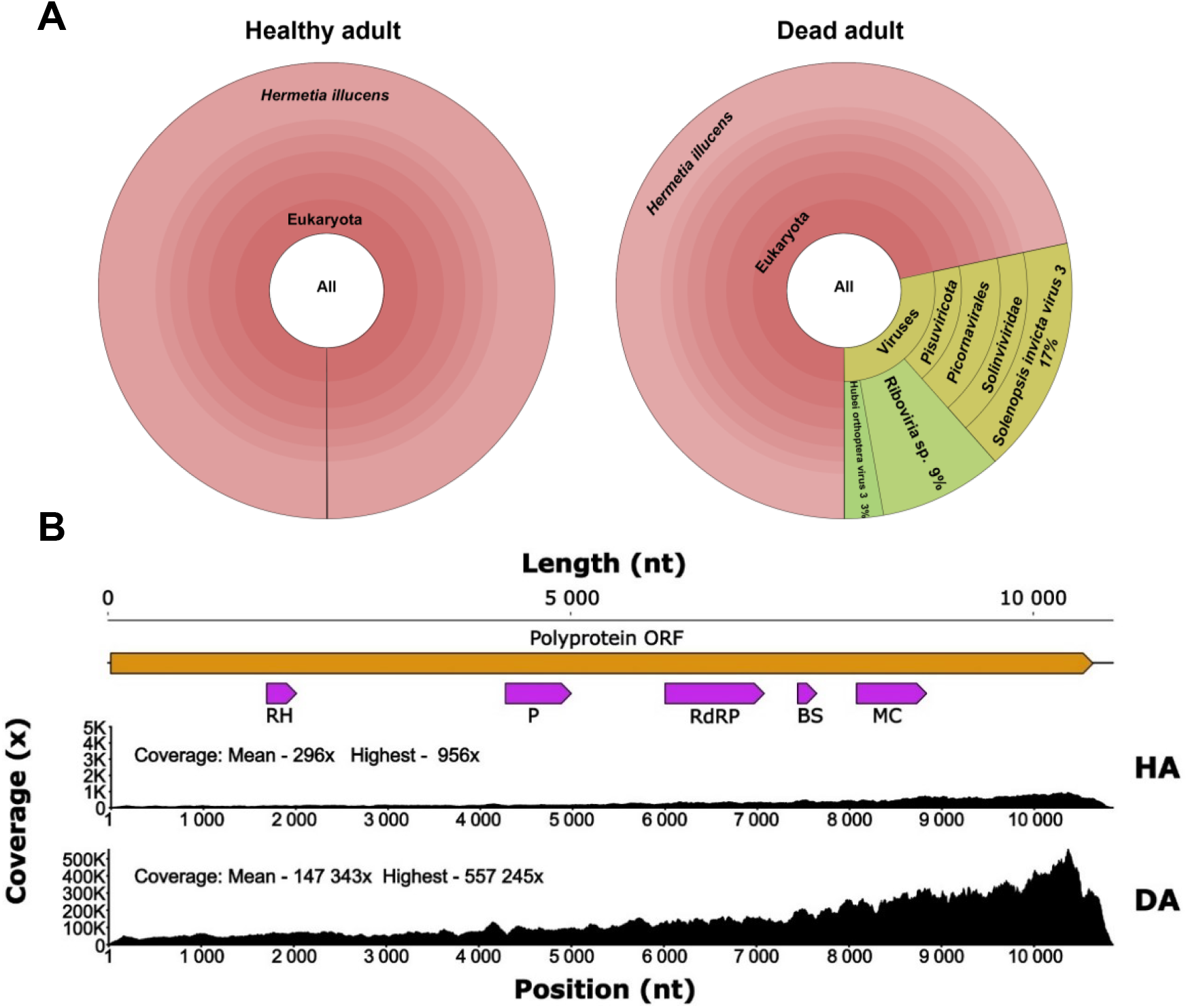
Detection of viral sequences in the BSF samples. (A) Visualisation of read classifications based on result outputs generated by Lazypipe2 analysis. For both the healthy adult (HA) and dead adult (DA), the minor taxa classifications (encompassing mostly bacteria and archaea) totalled less than 0.03% of transcripts. (B) Mapping of sequencing reads to the HiSvV genome. Annotated polyprotein ORF (orange) with putative conserved regions of proteins (purple) consist of an RNA helicase (RH), protease (P), RNA-dependent RNA polymerase 1 (RdRP), dsRNA binding site (BS) and a major capsid (MC). The upper coverage panel displays the HiSvV coverage in the healthy adult (HA), and the lower coverage panel shows the coverage of HiSvV in the dead adult (DA). Note the difference in Y-axis scales for the HA and DA samples.

### Icosahedral-like viral particles were identified in HiSvV positive samples

To confirm that HiSvV sequences derived from viral particles infectious to BSF and further characterize its viral structure, transmission and pathogenicity, we partially purified the viral particles. Homogenates (FH) from infected adults were processed to discard cellular debris and generate a partially purified (PPV) sample. According to the viral quantification, the purification procedure enriched the sample in about 4.3-fold with an increase in the relative abundance of HiSvV from 8 679 to 37 152 (Figure 2A). Given the prepurification protocol aims at selecting viral particles, finding an enrichment of HiSvV genomic equivalents indicated the viral nature of HiSvV.

**Figure 2.**
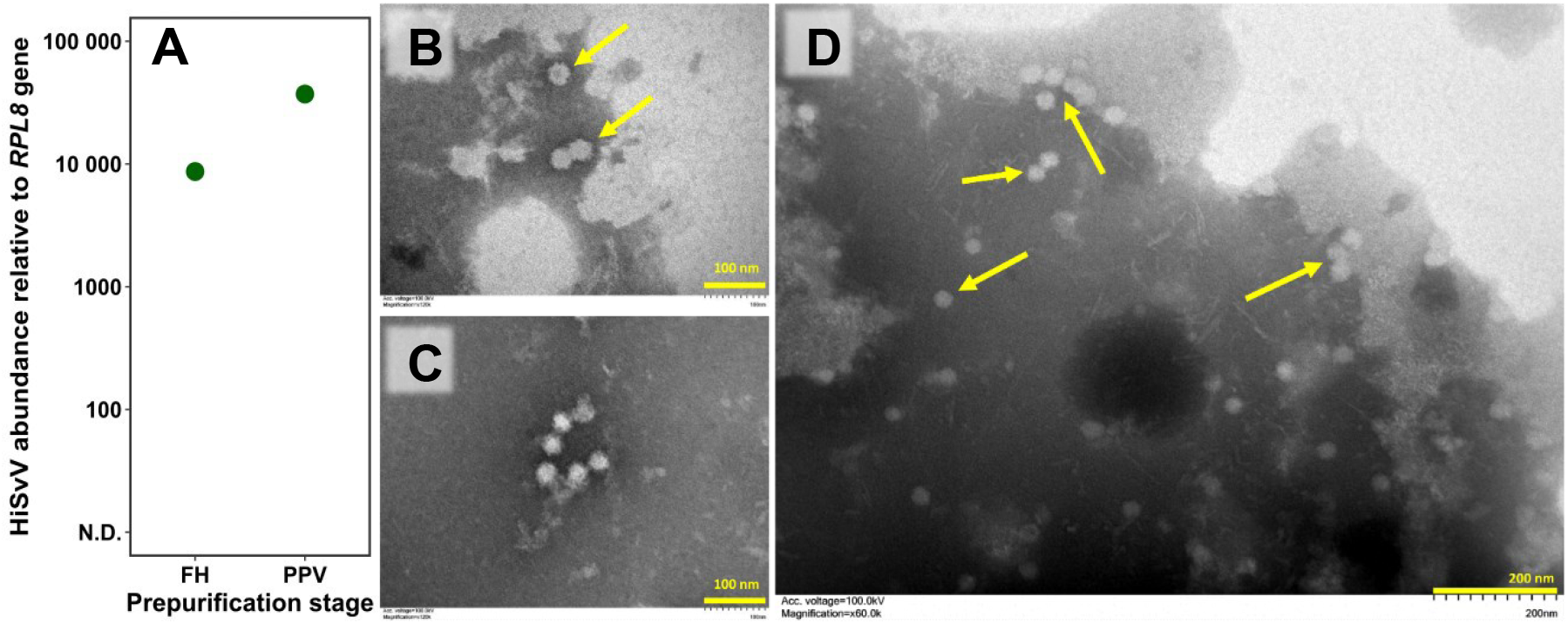
HiSvV purification and visualization by TEM. (A) Enrichment of HiSvV genomes after virus partial purification. Stages of the prepurification protocol compared were initial homogenate filtered through gauze (FH) and partially purified virus (PPV). (B to D) TEM observation of HiSvV particles in the PPV samples. The scale bars in the bottom right corner coloured in yellow represent 100 nm (B and C) and 200 nm (D).

Further exploration of the prepurified viral sample using TEM (Figure 2B-D) led to the visualisation of particles resembling in size and structure of other members of the *Solinviviridae* family [25]. These particles were observed isolated or aggregating together in a chain-like fashion and presented a similar morphology to other insect-infecting solinviviruses [25,27,28,68]. The capsid structure appears to be spherical with an icosahedral-like shape and a mean diameter of 33.9 ± 3 nm (Figure 2B-D).

### HiSvV replicates in BSF and has a systemic tropism

Initial trials to orally infect BSF larvae with PPV were unsuccessful and HiSvV was not detected in adults derived from two and seven days-old larvae exposed to concentrations of 6 x 10^9^ and 1.5 x 10^8^ HiSvV genomic equivalents/g of diet, respectively. In an alternative approach, BSFs in the prepupal stage were injected with the virus, and the viral abundance and tissue tropism was subsequently determined by RT-qPCR in adults which emerged on average 17 days after injection (Figure 3A and B). While HiSvV inoculation did not clearly impact adult emergence, HiSvV was detected consistently across the lifespan of both male and female adults (Figure S2; Figure 3A and B). Despite this, there was no drastic increase in the level of HiSvV genetic material within 13 days post-emergence (dpe) (Figure 3A). Mock-infected adults showed no HiSvV presence whether sampled within 24 hours or five dpe, confirming the specificity of the viral detection (Figure 3A). Following these observations, adults were dissected at three dpe to assess HiSvV tropism in various tissues (Figure 3B). Although HiSvV was found in all examined tissues, it was less prevalent in the reproductive tracts (including testis and ovaries). These results indicate a systemic infection, consistent as found in other *Solinviviridae* [27,68].

**Figure 3.**
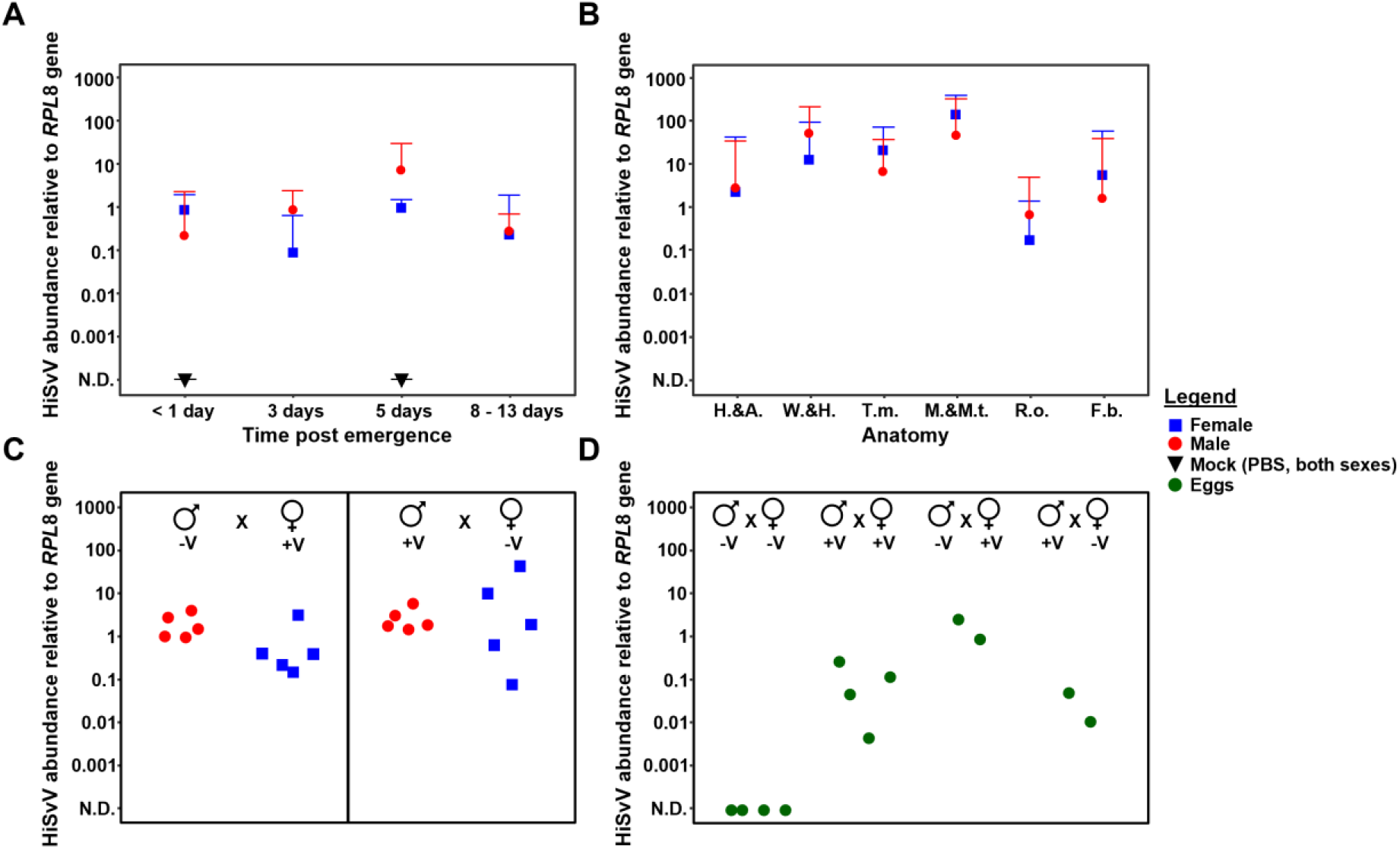
Detection of HiSvV in adults and egg clusters after prepupal injection. (A) Viral relative abundance at different days after emergence from pupae and (B) detection of HiSvV in different tissues of adults processed three days post emergence. The lower threshold for detection of HiSvV in samples was indicated by “N.D.”. Individuals injected with PBS were used as negative controls during inoculation. The error bars represent standard deviation (SD) of the mean on a log scale. Only the upper bars and caps of the SD are shown. (C & D) Testing horizontal and vertical transmission of HiSvV between BSF. Adults and eggs were obtained during cohabitation experiments and the level of HiSvV was assessed in the (C) adults and (D) egg clusters. Single male and female individuals were each represented by red dots and blue squares respectively. The green points represent single egg clusters tested. Individuals inoculated with HiSvV during prepupal stage are indicated using “V+”, while “V-” specifies mock-infected adults. “N.D.” on the y-axis indicates that HiSvV genetic material was not detected during qPCR.

### HiSvV is horizontally transmitted between adults and likely vertically transmitted

To determine HiSvV transmission routes, cohabitation experiments were performed using adults emerged after HiSvV injection (Figure 3C and D). Infected males were reared with non-infected females and *vice versa*. The relative abundance of HiSvV was analysed in the post-mated adult cadavers collected between 14 and 19 dpe, as well as in eggs from these crosses.

All tested adult cadavers from the cohabitation experiments were found positive for HiSvV, independently of their sex and their original infection status (Figure 3C). After cohabitation, viral titres were similar between the originally infected and non-infected adults. Moreover, HiSvV was detected in egg clusters where both parents were infected with HiSvV before cohabitation, and in egg clusters where at least one parent had been infected with HiSvV before cohabitation, independently of the infected sex (Figure 3D). HiSvV was not detected in the egg clusters from mock-infected parents (Figure 3D). These cohabitation results indicated HiSvV is horizontally transmitted between adult BSFs. In addition, since HiSvV was detected in egg clusters, the results also suggested probable maternal as well as paternal vertical transmission.

### HiSvV negatively impacts BSF adult survival after oral infection

Since premature mortality of BSF adults was one of the symptoms originally observed, the pathogenic nature of HiSvV was assessed by monitoring the survival of adults orally infected with HiSvV (Figure 4). Adults were orally inoculated 1 dpe to assess the effect on their lifespan. The relative abundance of HiSvV in both males and females increased similarly by almost four orders of magnitude and plateauing by 7 days post inoculation (dpi) (Figure 4A), confirming the successful oral infection of the adults. Focusing on adult survival, independent of the infective status, females exhibited shorter lifespans than males by 8 days on average (Figure 4B and C; Table S1 and S2). However, non-infected females were already exhibiting a hazard ratio of 6.5 (*p* < 0.001) when compared to the lifespan of non-infected males (Table S1A). Despite this, the hazard ratios of infections for both concentrations of HiSvV ranged between 3.5 to 4.9 (*p* < 0.001) when separating by sex, showing similar levels of effect between the two concentrations used (Figure 4B and C; Tables S1B). Overall, survival analyses also showed that for both males and females, HiSvV infection increased the probability of early death by 3.9 to 4.1 times (*p* < 0.001) (Figure 4B and C; Table S2).

**Figure 4.**
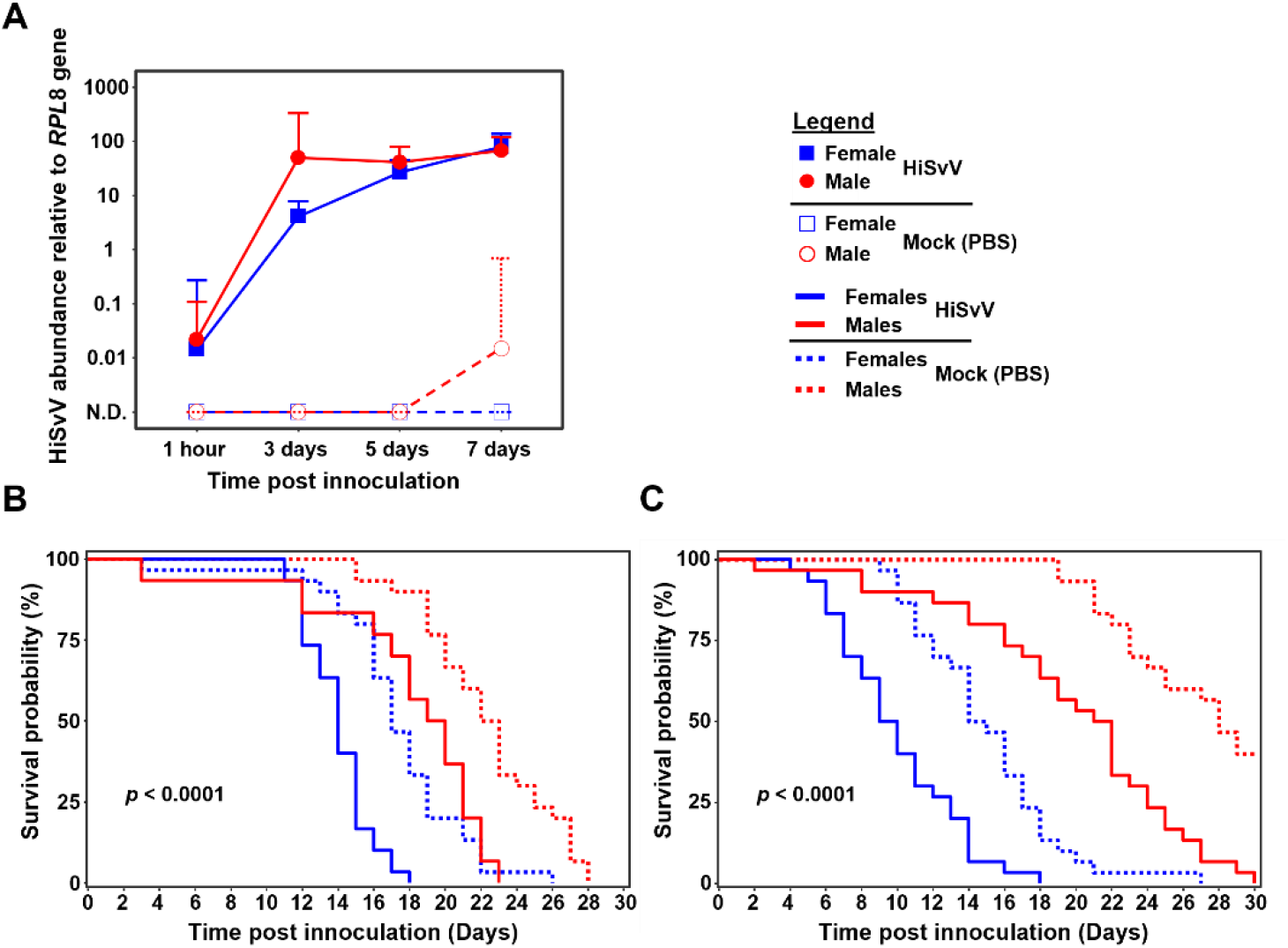
The effect of HiSvV oral infection on the survival of adult BSF. (A) Abundance of HiSvV in orally infected adults. Mean relative viral abundance in adults over seven days post-infection. Different colours represent the females (blue) and males (red). The filled shapes and empty shapes are used for the HiSvV- and mock-infected individuals, respectively. Lines were also used to indicate HiSvV (solid) and mock infections (dotted). The upper error bars represent the standard deviation (SD) of the mean on a log scale. (B & C) Impact of HiSvV infection on adult lifespan. Adults were orally infected with HiSvV PPV at two concentrations: 1.6×10^4^ genome equivalents/µl for experiment 1 (B) and 1.6×10^1^ genome equivalents/µl for experiment 2 (C). Males and females are differentiated by the colours red and blue respectively and a solid line separate HiSvV-infected from mock (PBS)-infected. The *p*-value within each plot was established by global comparison of strata within each experiment, indicating a statistical difference between all strata.

### HiSvV infection triggered a broad immune response in male BSFs

To determine host response to HiSvV infection, differential gene expression analyses were carried out on pooled males and females collected at 5 and 7 dpi. As similar viral titres and gene expression profile were obtained for 5 and 7 dpi samples, they were combined in subsequent analyses. Overall, differentially expressed gene analysis showed a broad response to HiSvV infection, with more genes being upregulated (UR) than downregulated (DR) in HiSvV infected adults (Figure 5). Initially, comparing significantly differentially expressed genes (DEGs) (*p*Adj < 0.05) in infected adults regardless of sex presented 799 UR genes and 486 DR genes (Figure 5A). However, when separating the males and females, the HiSvV-infected males showed a stronger level of DEGs (about 900) when compared with females (about 250) (Figure 5, Tables S3 and S4).

**Figure 5.**
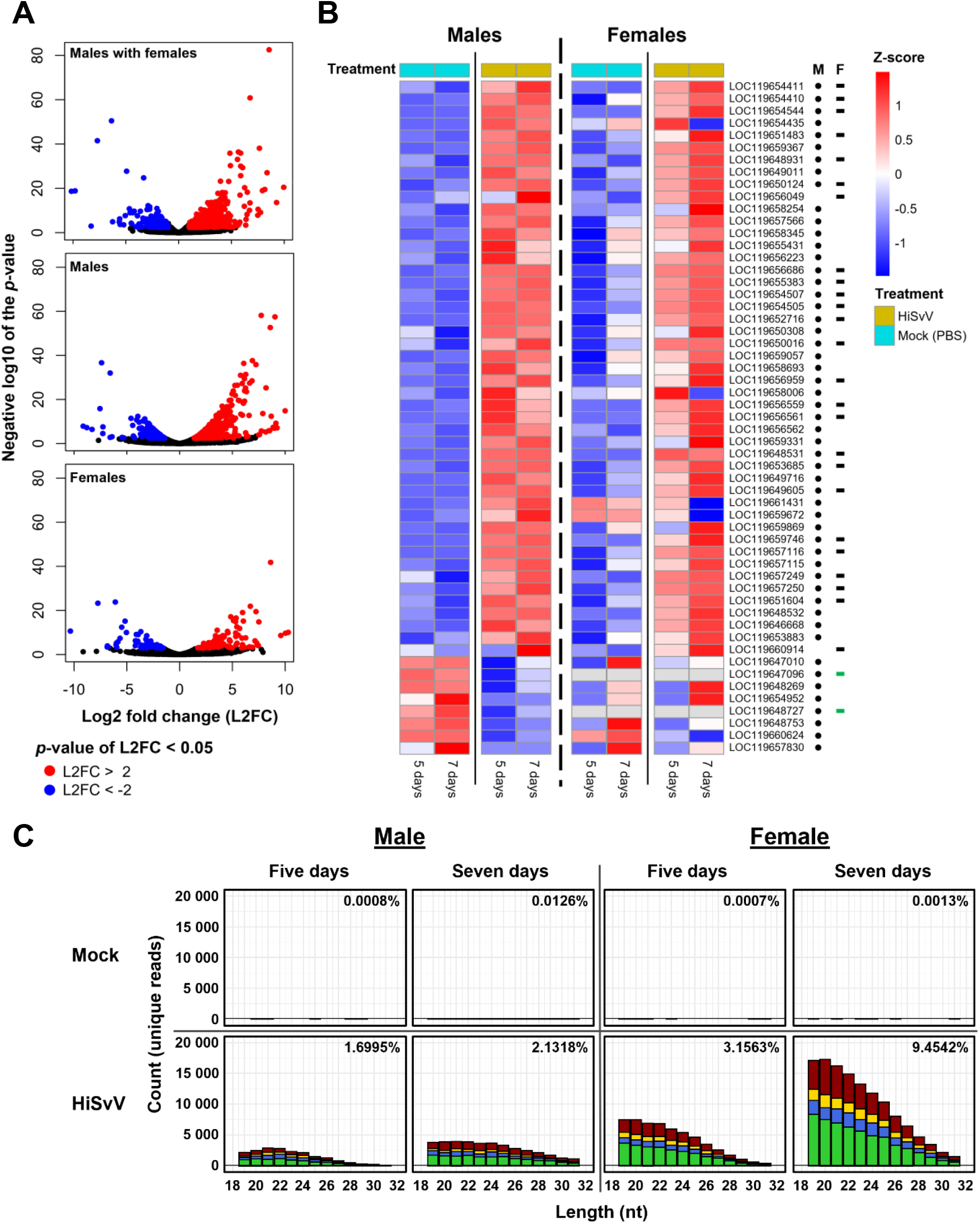
Analysis of transcriptional response to HiSvV infection in BSF adults. (A) Volcano plots depicting DEGs with negative and positive log2 fold-changes (L2FCs). (B) Heatmap of immune-related genes with a significant L2FCs contrasting HiSvV-vs Mock (PBS)-infected males and females. Males and females underwent DEQseq2 analysis separately for heatmap visualisation. The black dots represent genes with significant a L2FC in males the black dashes for females. The green dashes indicate genes where the gene count was “0”. Sizing small RNA from of HiSvV genome degradation. (C) Small RNA reads were mapped to the HiSvV genome in HiSvV-inoculated and mock-inoculated BSF. Profiles were obtained for adults collected at 5- and 7-days post-infection and in males and females. Small RNA sequences which started with an adenine were coloured in green, uracil in blue, thymine in red, guanine in yellow and ambiguous “N” in black. The percentage of mapped sRNA reads to the total number of reads in each sample was indicated in the top right corner of each subplot the bars above zero represent positive sense RNA mapping and the bars below represent negative sense mapping.

Although females had 8.5x less independent DEGs than males, males and females still shared 128 UR and 27 DR genes (Figure 5, Tables S3 and S4).

Closer examination of the DEGs found that 26.5% male DEGs and 31.4% of the female DEGs were uncharacterised (Tables S3 and S4). Analysis of biological processes through gene ontology (GO) enrichment pointed to at least 10 genes from the “response to viruses” process (FDR = 3.9e-04) including 8 involved in “defence responses to viruses” (FDR = 6.8e-04) in females. Additionally, 6 of these 10 genes were involved in the “negative regulation of viral genome replication” (FDR = 3.9e-04) (Figure S3). For males, there were at least 17 genes involved in “responses to viruses” (FDR = 8.8e-05), including 14 involved in “defence response against viruses” (FDR = 4.8e-04) and 9 involved in the “negative regulation of viral genome replication” (FDR = 4.2e-05) (Figures S3 and S4; Tables S3 and S4).

A deeper screening of biological processes from the same GO enrichment analysis as well as text and reference mining of male and female datasets found that at least 55 DEGs were likely related to immunity (Table 1; Figures S3 and S4; Tables S3 and S4). The vast majority of these genes (45 genes in males and 25 in females) were upregulated, suggesting the triggering of immune-related pathway activations after the infection (Figure 5B, Table 1). Of these 55 DEGs, most were linked to immune-related processes such as the Toll, IMD, and apoptosis pathways, with several encoding antimicrobial peptides (AMPs) including attacins and holotricins (Table 1). Infected flies showed sex-specific AMP regulation: two attacin-B-like genes were strongly upregulated in males and females, whereas holotricin-3-like genes displayed opposite regulation between sexes (Figure 5B). Males exhibited a broader overall immune response, while the putatively immune-related procathepsin L-like was uniquely expressed in females. In addition, several DEGs were associated with apoptotic and autophagy-related processes, as well as interferon-like responses, suggesting that programmed cell death and other antiviral mechanisms may contribute to the defence against HiSvV (Table 1). Only a few DEGs were associated with RNAi pathway, including argonaute-2-like and dicer, questioning the potential influence of RNAi mechanism against HiSvV infection (Table 1).

**Table 1.**
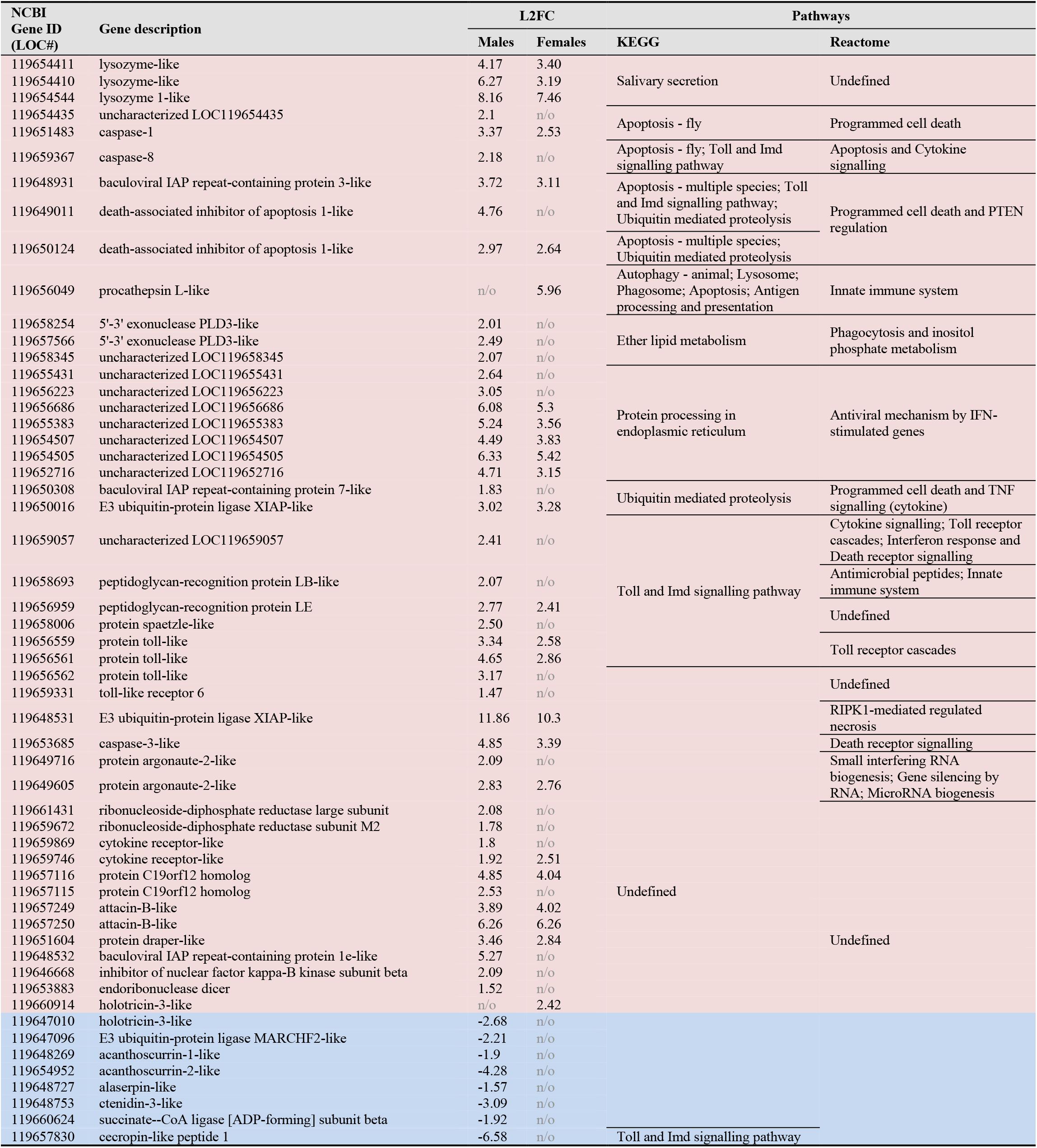
Putative Reactome and KEGG pathway annotations of significant (*p*Adj < 0.05) DEGs related to immune-linked biological processes. Both KEGG and Reactome annotations were statistically significant (*p* < 0.05). Genes coloured in red had an L2FC above 0, and those in blue were below 0. “n/o” indicates no significant L2FC observed.

### Absence of siRNA response to HiSvV infection in adult BSFs

Although only argonaute-2 and endoribonuclease dicer potentially involved in the RNAi defence pathway were found to be regulated, we decided to explore if this defence mechanism was triggered by HiSvV infection. Small RNA (sRNA) profiles were thus examined after HiSvV infection (Figure 5C). Mapping of sRNA reads to the HiSvV genome showed an increase in counts and percentage of mapped reads from 5 to 7 dpi (Figure 5C), further supporting active viral infection as detected earlier (FIGs 3A and 4A). However, the broad size range from 19 to 31 nt, targeting the HiSvV viral genome without a prominent peak at 21 nt suggested that the typical siRNA antiviral pathway was not activated (Figure 5C). The widespread distribution of the sRNA profile observed rather indicated the degradation of HiSvV genomic material and suggested an alternative defence response.

## Discussion

Pathology research in mass-reared insects for food and feed has garnered more attention within the last few years, and BSFs are no exception [1,4,6,16,69]. While advances in virus research have been made in other mass reared insects such as crickets and mealworms, they are just developing in BSFs [4,16,19,70,71]. Our study was the first to isolate a virus from diseased BSFs and to demonstrate its pathogenicity in this species, a notable finding given the economic importance of BSFs [2,72].

We received BSF adult samples from a rearing facility experiencing high levels of adult mortality and reduced egg production. The presence of HiSvV genetic material was linked with the presence of viral particles, confirming it as a viral agent with a morphology consistent with members of the *Solinviviridae* family [25]. Horizontal transmission of HiSvV was demonstrated through successful experimental oral infection of BSF adults as well as through cohabitation experiments in which HiSvV was detected in exposed adults and in the egg clusters. Although the presence of HiSvV in egg clusters suggest HiSvV potential for vertical transmission, the systemic tropism favouring the digestive tract indicates that the typical infection occurs mainly through the oral-faecal route. This was further confirmed by oral inoculation of adults as also found for a closely related solinvivirus, SINV3 [33,73]. Whether HiSvV can sustainably replicate in larvae remains unresolved, as for other solinviviruses [27,28,37].

HiSvV infection in adults reduced the lifespan of both males and females. This is the first study examining the effect of a solinvivirus infection on sexes, contrasting prior studies performed on unsexed larvae of shrimp or on female castes and larvae in ants and bees [28–31]. Other *Solinviviridae*, such as SINV3, AmSV1 and PvSV, also cause premature mortality that have been linked to colony collapse in ants, honeybees, and whiteleg shrimp [29–31]. In addition, the risk of colony collapse due to HiSvV infection may be reinforced if BSF are dying before mating or oviposition, particularly if virus levels are allowed to build up within the rearing facility [16]. This risk would be more pertinent for continuous rearing setups, as mating can start at least three to four days post-emergence [76,77].

To better understand host-pathogen interactions and potential sex-specific effects, we explored the immune response in both males and females using DEG analysis and sRNA profiling. The DEG analysis revealed significant transcriptional changes in antiviral immunity-related genes (Figures S2 and S3; Tables S4 and S5), indicating a clear immune-related response to an entomopathogenic virus introduced at a relatively low concentration. The sRNA profiles from HiSvV-infected flies were broad rather than displaying specific peaks, suggesting that BSFs did not mount a specific silencing response against HiSvV. This pattern implies that the virus may evade sRNA pathways, a strategy known in other insect-infecting *Solinviviridae* and *Dicistroviridae*, which can avoid RNAi silencing through a dsRNA binding domain on their genomes [28,78–80]. *Hermetia illucens* solinvivirus also encodes for a homolog to this dsRNA binding domain, supporting the interpretation that the broad sRNA profiles reflect general viral genetic material degradation rather than specific RNAi or PIWI responses. Males exhibited a broader immune response than females, and they generally survived longer, suggesting that a major activation of the immune defences may contribute to reduce the detrimental effects of the viral infection. Similarly, differences in immune activity between sexes has been observed in other flies, e.g. *Drosophila* species, reinforcing the need to examine impacts of BSF viruses in both sexes when possible [79,81–85].

It remained to be determined whether the upregulated antimicrobial peptides (AMPs) in our study were directly involved in antiviral defence or instead target opportunistic microbiota [12,28,79,80]. While not all AMPs act against viruses, some cecropins, attacins, and defensins are known to participate in antiviral responses [79,86]. Both sexes showed significantly higher L2FC values for attacin-B-like genes. Holotricin-3 which has antimicrobial properties against bacteria and fungi but unclear antiviral activity [87,88], was downregulated in males yet

upregulated in females, possibly reflecting a sex-specific role in HiSvV defence. The upregulation of genes linked to recognition, signalling, and other processes in the IMD and Toll pathways, further supports a putative antiviral role for effectors linked to these pathways such as attacin-B-like and holotricin-3-like (putative defensin) in BSF-HiSvV interactions [79,87,89]. The apparent number of genes associated with programmed cell death, autophagy, and interferon-like responses suggests these processes may play a role in degrading HiSvV-infected cells, which could explain the broad, degraded sRNA profiles. These responses are defence mechanisms likely effective against pathogens that evade RNAi and AMPs and are particularly relevant for insect-infecting picornaviruses such as *Dicistroviridae* and *Solinviviridae* [28,78–80].

Despite the “popular belief” that BSFs were insensitive to viral infections, this study showed for the first time that a recently discovered virus, HiSvV, can significantly induce premature mortality of BSF adults and is likely the culprit behind a reported loss of production within a BSF farm. Given the capacity of HiSvV to decrease production, this prompts the need to develop management plans and surveillance tools to prevent future outbreaks and interfacility transfer. Lastly, this work represents an important milestone in BSF pathology since it is the first study to show that viruses could be a threat to BSF rearing facilities.

## Acknowledgements

We would like to acknowledge the assistance by members of the INSECT DOCTORS consortium, IRBI and BIOTECMED institutes for their assistance throughout the study. Namely the assistance by Corentin Clavé, Alexandra Cerqueira de Araujo, Thibaut Josse, Carole Labrousse, Melissa Lloyd, Marina Millet, Carlos Lopez-Vaamonde, Perrine Lutanie, Ariel Muñez Sánchez and Valentin Pressoir during experiments. We would also like to thank Alejandro Torres Ruiz from Entomotech S.L (Almeria, Spain) for providing the BSF seed colony to be used in the experimentation. This research was supported by the INSECT DOCTORS program, funded under the European Union Horizon 2020 Framework Programme for Research and Innovation (Marie Sklodowska-Curie Grant agreement 859850). Salvador Herrero’s research on BSF is also supported by grants TED2021-130679B-I00 funded by MCIN/AEI/10.13039/501100011033 and by the “European Union NextGenerationEU/PRTR”. Finally, we would like to thank the Servicio Central de Soporte a la Investigación Experimental (SCSIE) for the use of equipment and assistant for performing the transmission electron microscopy.

## Supplementary data

**Figure S1.**
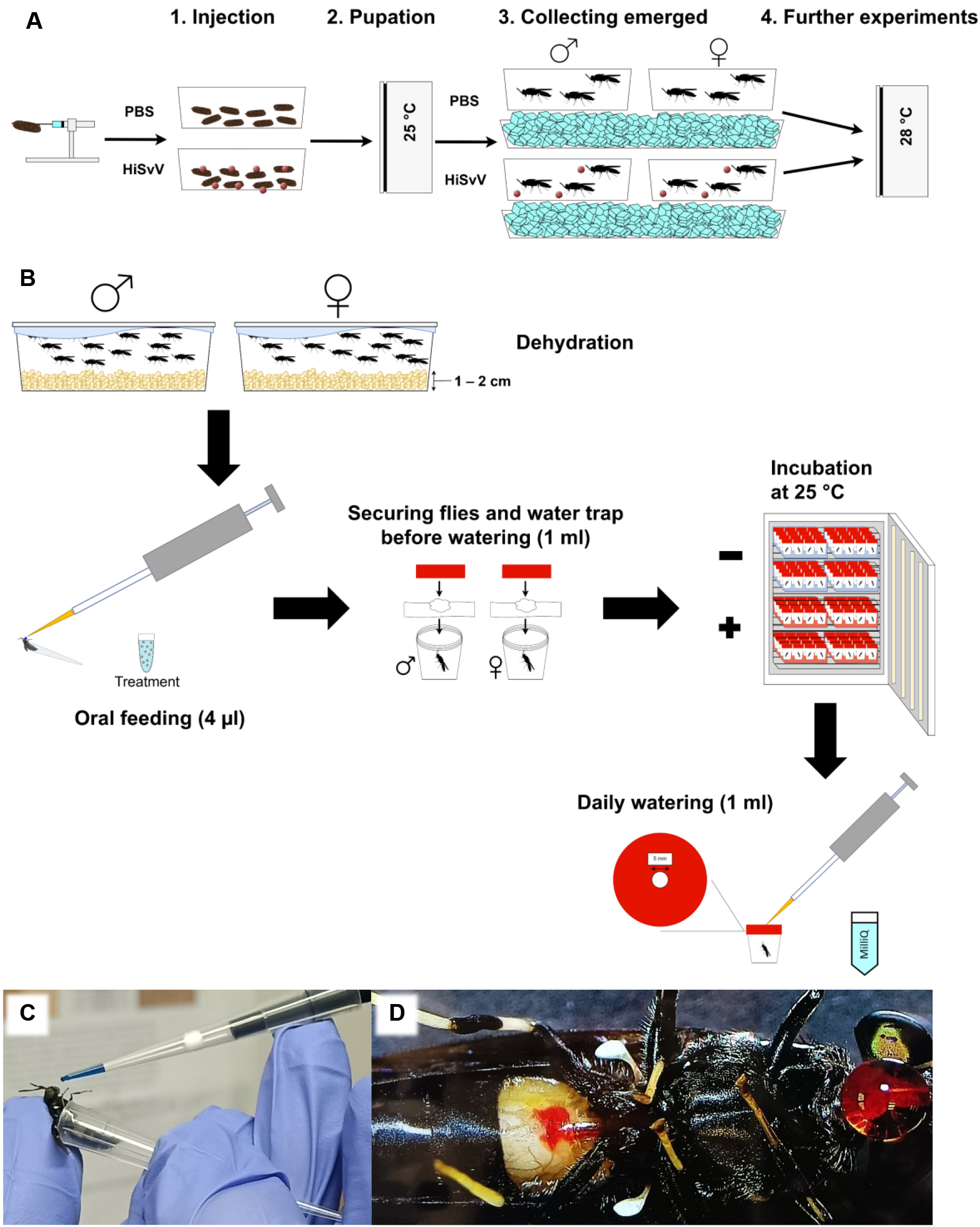
Injection and oral bioassay protocols used to study viruses in BSF. (A) Outline of HiSvV inoculation by injection of prepupae. After injection, prepupae were pooled by treatment and pupated at 25 °C. Emerged adults were sexed on ice and separated incubated at 28 °C. B) Visualisation of the droplet feeding bioassay starting with the dehydration process and then oral feeding by pipette, placing the dosed flies into cups and providing water, followed by the incubation period and daily watering. C) An adult held steady in a P20 pipette tip while receiving a droplet of virus inoculum. D) Red dye visible in the midgut of an adult after ingestion of solution. The experiments were performed with either a red or blue food colouring.

**Figure S2.**
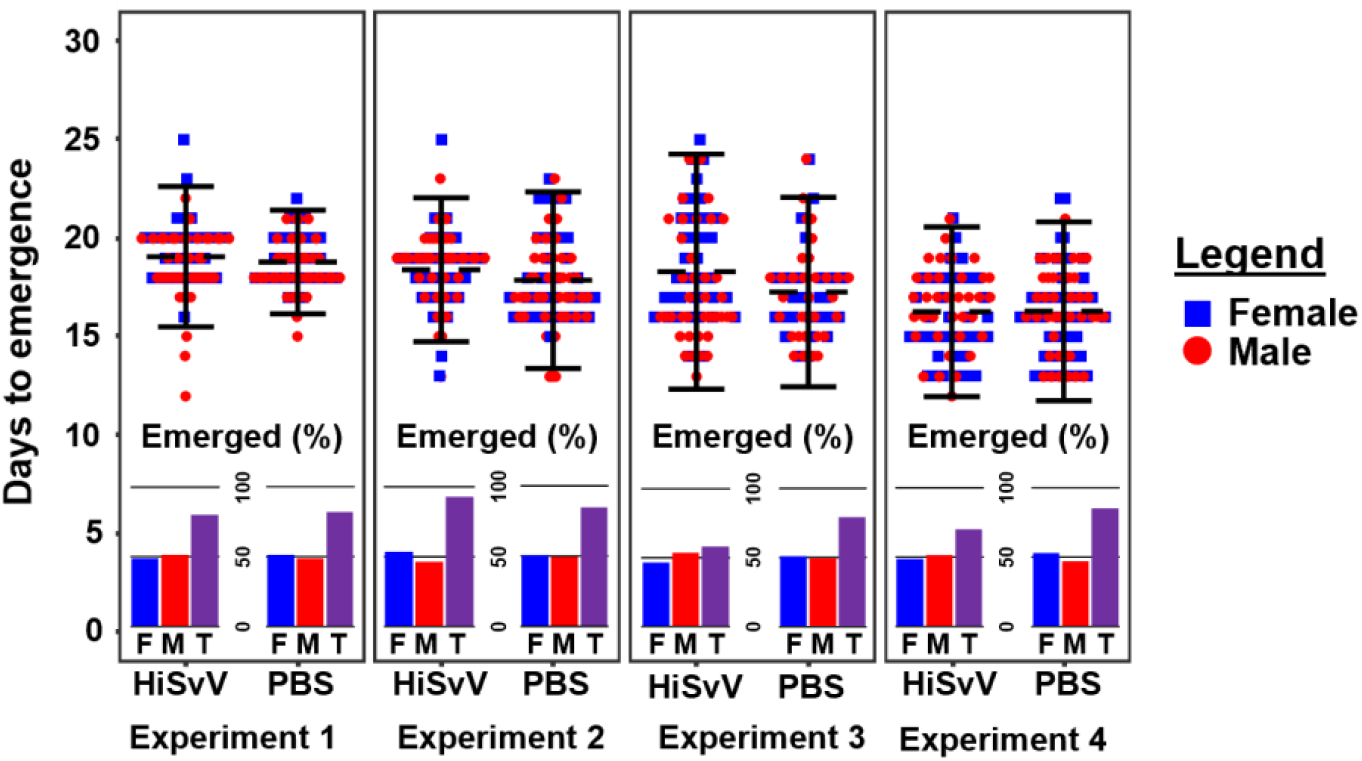
Adult emergence following prepupal injection-based inoculation of HiSvV. Time and emergence rate after viral infection (by injecting prepupal stages). Results correspond to four independent experiments. Each dot corresponds to a single adult. The sex of the obtained adults is indicated with different colours. The error bars indicate the standard deviation of the mean within each treatment group. Bar at the bottom the plot depicted the percentage of emerged males (M, red), females (F, blue) out of the total percentage of emerged adults (T, purple).

**Table S1.** Statistical analyses models inferred from survival of adult BSFs over 30 days when infected with HiSvV.

| Analysis of Deviance Table (Type II tests) |  |  |  |  |
| --- | --- | --- | --- | --- |
| LR | Chi<sup>2</sup> ($\chi^2$) | Df | p-value | |
| Sex | 66.94 | 1 | 2.80e-16 | *** |
| Experiment | 1.88 | 1 | 1.70e-01 |  |
| Sex:Experiment | 20.31 | 1 | 6.59e-06 | * |

| Hazard ratio of coxph model: Surv(death, status) ~ sex + experiment |  |  |  |  |  |
| --- | --- | --- | --- | --- | --- |
|  | Coef. | HR | SE (coef.) | z | p-value |
| Females | 1.87 | 6.48 | 0.23 | 8.07 | 7.02e-16 *** |
| Experiment 1 | 0.29 | 1.33 | 0.21 | 1.36 | 1.73e-01 |

**B**
| Hazard ratio of coxph model: Surv(death, status) ~ treatment |  |  |  |  |  |  |
| --- | --- | --- | --- | --- | --- | --- |
| Sex | Dose (GE/μl) | Coef. | HR | SE (coef.) | z | p-value |
| Males | 1.6x10<sup>4</sup> | 1.26 | 3.52 | 0.31 | 4.06 | 5.02e-05 *** |
|  | 1.6x10<sup>1</sup> | 1.43 | 4.19 | 0.31 | 4.57 | 4.96e-06 *** |
| Females | 1.6x10<sup>4</sup> | 1.59 | 4.89 | 0.32 | 4.96 | 6.89e-07 *** |
|  | 1.6x10<sup>1</sup> | 1.25 | 3.48 | 0.28 | 4.38 | 1.19e-05 *** |

**Table S2.**
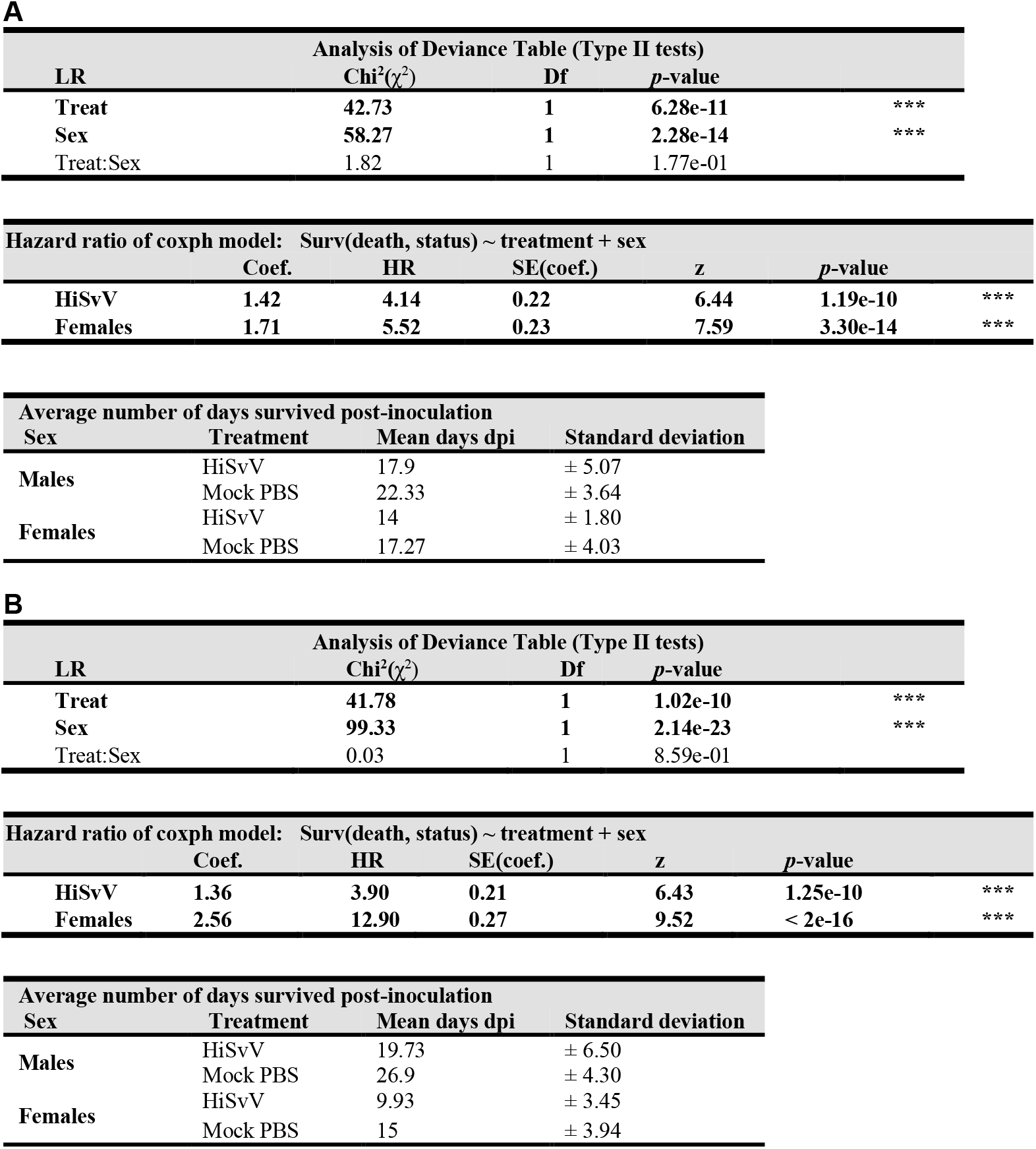
Hazard ratios and deviance-based analysis of HiSvV infection in adult BSF. Adults were infected with two concentrations of HiSvV PPV: 1.6×10^4^ genome equivalents/µl (A) and 1.6×10^1^ genome equivalents/µl (B). **A**

**Figure S3.**
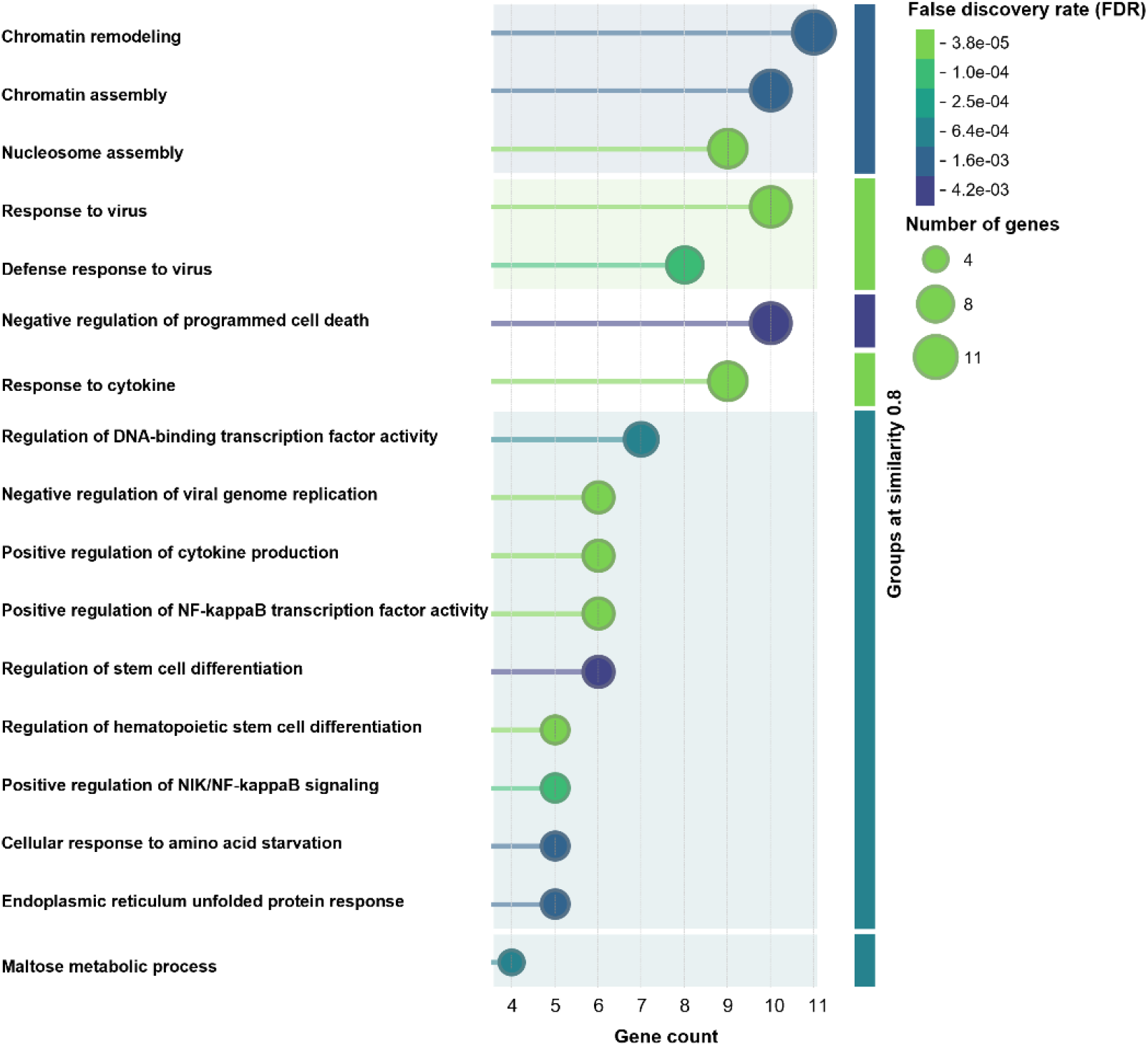
Putative biological processes of significant DEGs in BSF females infected with HiSvV.

**Figure S4.**
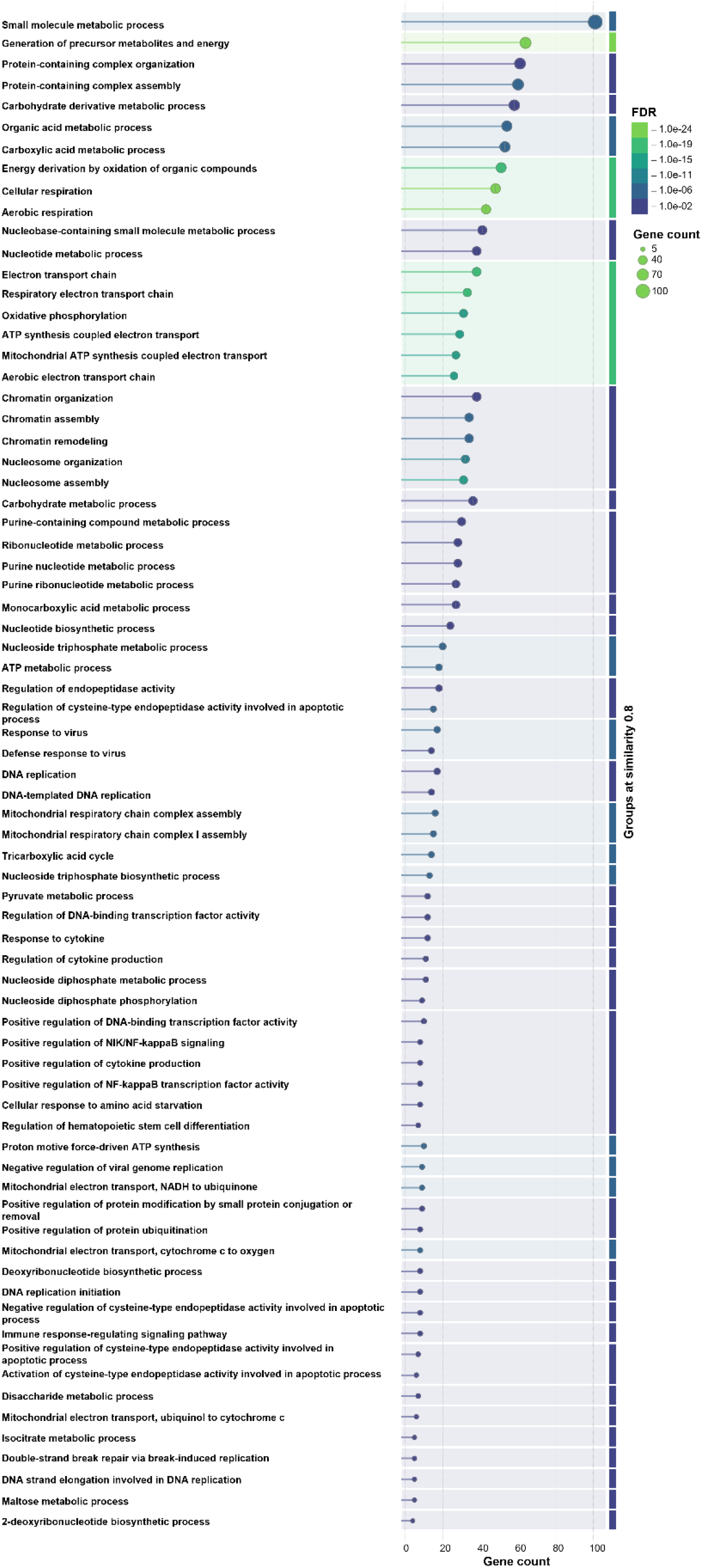
Putative biological processes of significant DEGs BSF males infected with HiSvV.

